# The subesophageal ganglion modulates locust inter-leg sensory-motor interactions via contralateral pathways

**DOI:** 10.1101/261164

**Authors:** Daniel Knebel, Johanna Wörner, Jan Rillich, Leonard Nadler, Amir Ayali, Einat Couzin-Fuchs

**Author notes:** Equal contribution (Daniel Knebel, Johanna Wörner). Author for correspondence: Einat Couzin-Fuchs.

## Abstract

The neural control of insect locomotion is distributed among various body segments. Local pattern-generating circuits at the thoracic ganglia interact with incoming sensory signals and central descending commands from the head ganglia. The evidence from different insect preparations suggests that the subesophageal ganglion (SEG) may play an important role in locomotion-related tasks. In a previous study, we demonstrated that the locust SEG modulates the coupling pattern between segmental leg CPGs in the absence of sensory feedback. Here, we investigated its role in processing and transmitting sensory information to the leg motor centers, and mapped the major related neural pathways. Specifically, the intra- and inter-segmental transfer of leg-feedback were studied by simultaneously monitoring motor responses and descending signals from the SEG. Our findings reveal a crucial role of the SEG in the transfer of intersegmental, but not intrasegmental, signals. Additional lesion experiments, in which the intersegmental connectives were cut at different locations, together with double nerve staining, indicated that sensory signals are mainly transferred to the SEG via the connective contralateral to the stimulated leg. We therefore suggest that, similar to data reported for vertebrates, insect leg sensory-motor loops comprise contralateral ascending pathways to the head and ipsilateral descending ones.

## 1 Introduction

Animal locomotion is based on multidirectional, dynamic interactions within and between several control levels: central pattern-generating neuronal circuits, sensory inputs that modulate these circuits, and higher neuronal centers residing in the brain or head ganglia.

Ample research across several decades has focused on the central pattern generators (CPGs), demonstrating that these neuronal oscillators constitute the basis of practically all rhythmic behaviors, including locomotion (Marder and Bucher 2001; Mulloney and Smarandache 2010; Ayali and Lange 2010; Rybak et al. 2015). Similarly, extensive research has been dedicated to the role of sensory feedback in shaping the CPG-induced behavior in different animals. In legged-locomotion specifically, sensory inputs have been attributed a role in the functional adaptability of leg motor patterns to the changing environment (e.g., Altman 1982; Burrows 1987; Pearson 1993, 2004; Zehr and Stein 1999; Büschges and Pearson 2006; Ayali et al. 2015b). Leg proprioception during walking has been shown to be instrumental for inter-leg coordination in the stick insect (e.g. Cruse 1990; Büschges et al. 2008; Borgmann et al. 2009) and, similarly, mechano-sensory feedback was reported to mediate intersegmental information transfer and adjust locomotion patterns in cockroaches (Zill et al. 2009; Fuchs et al. 2012; Ayali et al. 2015a; Couzin-Fuchs et al. 2015a, b). The effect of local sensory input is context- and task-dependent, influenced by the activity of neighboring legs (Knop et al. 2001; Hellekes et al. 2012) and descending information (Ridgel et al. 2007; Mu and Ritzmann 2008).

The higher motor centers provide additional control of locomotion. The insect central complex, for example, was shown both to process environmental cues and to execute appropriate commands for walking, such as speed change and turning (for review: Pfeiffer and Homberg 2014). The subesophageal ganglion (SEG), one of the insect head ganglia, which is located below the esophagus (Ito et al. 2014). Beyond its role in local control of the mouthparts (Rand et al. 2008, 2012), the SEG was shown to play an important role in insect locomotion. Ablating the SEG in cockroaches was demonstrated to affect walking and initiation and duration of escape (Gal and Libersat 2006; Kaiser and Libersat 2015). Recordings from locust SEG descending interneurons (DINs) demonstrated elevated activity during and after the preparatory phase of walking (Kien 1990a). These DINs typically fire throughout the walking bout, as temporally-structured patterns that are not directly correlated with the stepping cycles (Kien 1990b). It has recently been shown in *Drosophila* larvae that a small group of neurons in the subesophageal zone contribute to the control of larval chemotaxis (Tastekin et al. 2015). Inhibiting these neurons compromised the timing and coordination of reorientation maneuvers while activation of these neurons through optogenetics and thermogenetics were sufficient to initiate reorientation maneuvers. In *in-vitro* preparations, removal of the subesophageal zone affected the degree of asymmetry of fictive crawling motor patterns (Pulver et al. 2015). However, the intricate interactions among CPGs, sensory information, and higher motor centers, which together generate coordinated walking behavior, are still far from fully understood.

In an attempt to characterize the interplay among the different neuronal elements that control walking in the desert locust, *Schistocerca gregaria*, we previously mapped the central basis of inter-leg coordination in the thoracic ganglia (Knebel et al. 2017a). Following that, we demonstrated the role of the SEG in modulating the coupling between segmental leg CPGs (Knebel et al. 2017b). These findings are supported by anatomical descriptions showing that the SEG contains both descending and ascending interneurons and modulatory projections that reach all thoracic ganglia and the brain (Kien and Altman 1984; Bräunig 1991; Roth et al. 1994), and thus may be important in translating higher motor commands into direct execution signals.

In the current study, we revealed the role of the SEG in integrating proprioceptive information from the legs and generating intersegmental responses. To this end, we investigated the interactions between descending signals from the SEG and walking-related thoracic sensory-motor responses by means of extracellular recordings from nerves containing leg depressor and levator motoneurons during stimulation of sensory nerves, before and after lesioning the connections between the SEG and the thoracic ganglia. Our findings indicate a contribution of the SEG in intersegmental sensory-motor interactions.

## 2 Materials and Methods

### 2.1 Animals and preparation

All experiments were performed on adult desert locusts, *Schistocerca gregaria,* from our breeding colonies. Extracellular recordings were performed on *in-vitro* preparations of the thoracic ganglia and the SEG as follows: locusts were anesthetized with CO2, legs and wings were cut and the dorsal part of the head, including the supraesophageal ganglion (brain), was removed by a horizontal cut ventral to the compound eyes. After excising the abdomen, the thorax was opened along the dorsal midline. Gut, fat tissue, air sacs, and cuticular parts, including the tentorium from the head cuticle, were carefully removed, exposing the ventral nerve cord. All peripheral nerves were cut close to the ganglia, except for the meso- and metathoracic leg nerves 5 (numbered after Campbell, 1961), which were cut as distally as possible. Finally, the circumoesophageal connectives and the connectives between the last thoracic- and the first abdominal ganglia were cut, and the thoracic-SEG ganglia chain with its surrounding tracheal supply was dissected out of the body cavity, pinned in a clean Sylgard dish (Sylgard 182 silicon Elastomer, Dow Corning Corp., Midland, MI, USA), dorsal side up, and bathed in locust saline (in mM: 150 NaCl, 5 KCl, 5 CaCl2, 2 MgCI2, 10 Hepes, 25 sucrose at pH 7.4). The two main tracheae were opened and floated on the saline surface.

### 2.2 Electrophysiological recordings

Electrophysiological experiments were carried out at both Konstanz and Tel Aviv Universities. We used custom-made suction electrodes: a) to record extracellularly the activity of the 5A nerves, which contain three motor axons (the slow and fast trochanteral depressors and a common inhibitor); and b) to stimulate nerve 5B, which contains the majority of the leg sensory branches. This branch was chosen as a target for generating a general, unspecific, leg sensory stimulation. Thus, although this did not allow investigating very specific sensory modulations, it enabled us to trace general sensory-motor routes within the nervous system. To monitor activity from the connectives, we used silver wire hook electrodes. After dissection, and after every cut of the connectives, we left the preparation to recover for 15 minutes before recording.

Electrical stimulation of semi-intact preparation were programmed using a Grass S88 (Grass Instruments) and delivered by an electrode made out of 2 pins which was placed approximate to the N5b nerve in the hind leg. The stimulation protocol was composed of a pulse train (200 ms of 200Hz pulses of 1 ms each). To adjust the amplitude of these stimulations, the voltage was gradually increased till a response of the middle leg was apparent and then fixed to this amplitude for the rest experiment. The different stimulation properties needed to elicit a response are probably the result of the variance in the quality of the contact between the electrode and the nerve. To stimulate sensory nerves *in vitro*, either custom-made suction electrodes (in Konstanz University) or hook electrodes (in Tel Aviv University) were fashioned and used as described above. Stimulation protocols were either programed in Spike2 and delivered with Master 8 stimulator or programed and delivered with Master 8 stimulator to give pulse trains of 100-800ms of 100-200 Hz of 2 ms, with amplitude determined as above. The properties were fixed to the first response recorded, and remained consistent within each experiment, and were probably different due to the variance in conductance between the electrode and the nerve among the experiments. The N5b branch was chosen as a target for generating a general, unspecific, leg sensory stimulation. Thus, although this did not allow investigating very specific sensory modulations, it enabled the tracing of general sensory-motor routes within the nervous system.

### 2.3 Nerve staining

The neck connective was cut unilaterally and subsequently backfilled *in situ* with the tracer Neurobiotin (Vector Laboratories, Inc.). The specimen was placed in a moist chamber and stored for 24 h at 4°C to allow the tracer to diffuse along the cut neurons. Following 24 h of diffusion the interconnected suboesophageal, prothoracic, mesothoracic, and metathoracic ganglia were isolated together with the trachea system, which was opened at both ends and pinned on a Sylgard dish. The metathoracic 5B nerve was then backfilled with Dextran, Tetramethylrhodamine (Molecular Probes). Following an additional 24 h of diffusion at 4°C all remaining tracheas were removed from the ganglia. The nerve cord was then fixed for 2 h in 4% paraformaldehyde at room temperature. Subsequently, the specimen was washed 4 x 15 min in 0.1 M phosphate buffer (PB, pH 7.2), dehydrated in an ascending ethanol series (50%, 70%, 90%, 100%, 10 min each), washed 2 x 5 min in methyl salicylate (Merck KGaA), rehydrated in a descending ethanol series (100%, 90%, 70%, 50%, 10 min each) and washed again in 0.1 M PB (2 x 15 min).

To visualize the Neurobiotin trace the specimen was bathed in 0.1 M PB + 1% Triton X-100 (SigmaAldrich) and Streptavidin, Alexa Fluor 647 conjugate (Jackson ImmunoResearch Labs) in a lightproof box overnight at room temperature. The specimen was then washed 3 x 15 min in 0.1 M PB, dehydrated in an ascending ethanol series (50%, 70%, 90%, 100%, 10 min each), and finally cleared in methyl salicylate until full transparency. The whole-mount preparations were scanned with a laser scanning confocal microscope (LEICA TCS SP2, Leica) to capture the fluorescent signals emitted from Tetramethylrhodamine and Alexa Fluor 647. Resulting image stacks were processed with FIJI (http://fiji.sc/wiki/index.php/FIJI).

### 2.4 Data Analysis

Spike detection, analysis, and statistics were performed in Matlab (MathWorks, USA Inc.). Spike detection in the motor nerves was based on their amplitude, taking into account only excitatory motor units (slow and fast trochanteral depressors), without separating between them (see recording of nerve 5A in Figure 1a). In all analyses, we compared motor activity recorded 4 sec before stimulation with that recorded 4 sec after. To characterize correlation between motor units at the meso- and metathoracic ganglia, we transferred the spike timing data into vectors of activity density (with a sliding window of 100 msec bins). We used a 4 sec time window from the stimulation onset and calculated the Pearson correlation coefficient between the meso- and metathoracic activity vectors, and only statistically significant rho values were taken into account. Analysis of connective recordings was based on spike template recognition using Dataview software (University of St. Andrews), followed by overlay of the events in the different channels to identify descending signals, as further elaborated in Knebel et al. (2017b).

**Fig. 1.**
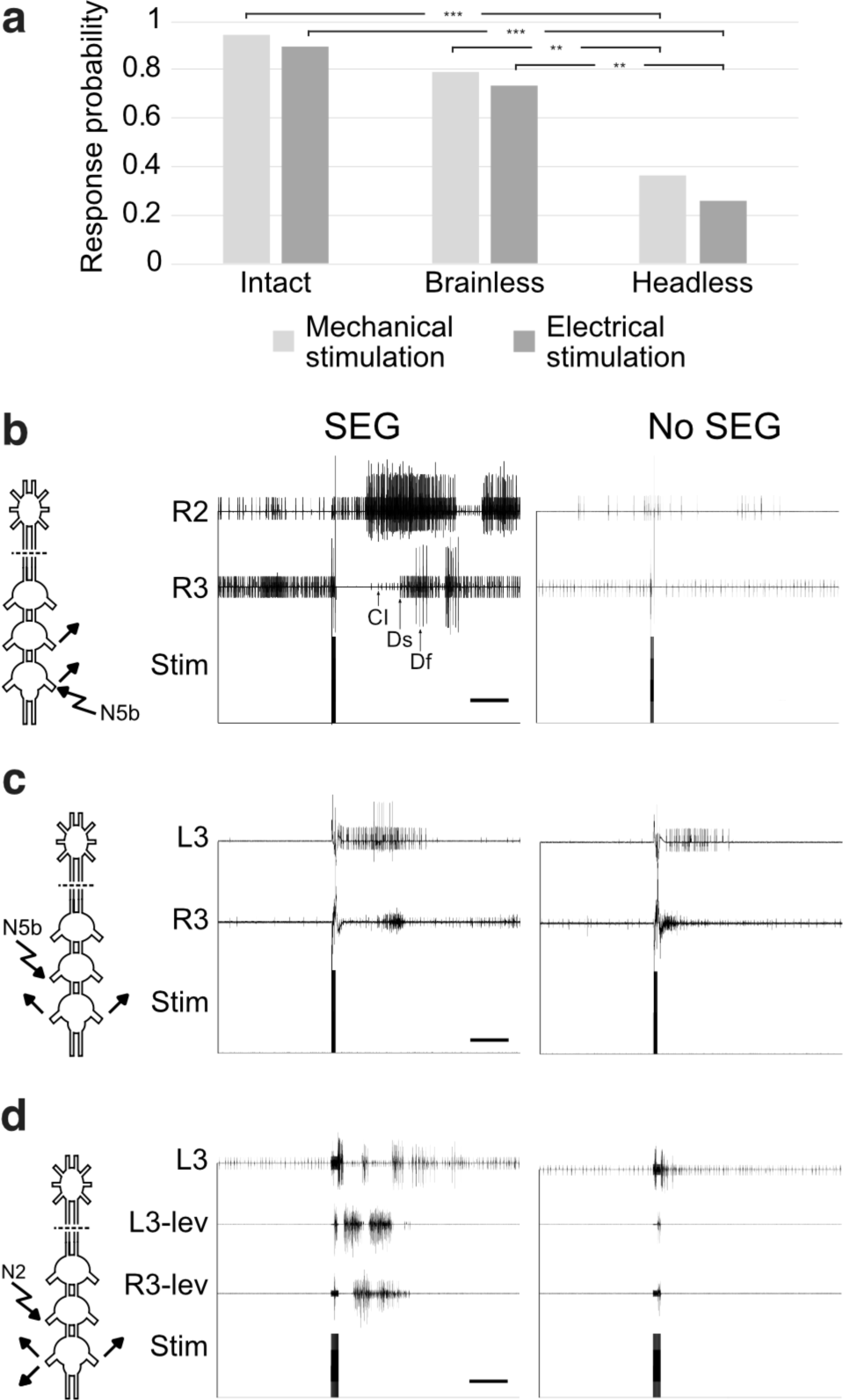
Responses to sensory stimulations, with and without the SEG. (A) Response probabilities of the middle leg to mechanical and nerve stimulation of the hind leg in intact, brainless and head less animals. **p<0.01 ***p<0.001. (B) Recordings from the ipsilateral meta- and mesothoracic depressors during stimulation of the metathoracic N5b, with and without the SEG intact (left and right panels, respectively). The different units in the depressor nerves are tagged (Df – fast depressor, Ds – slow depressor, CI – common inhibitor). Scale bar is 1 sec. (C) As in (B), but for stimulating the mesothoracic N5b and recording from the ipsi- and contra-lateral metathoracic depressors. (D) As in (B) but for recordings of the meta-thoracic levator and depressor nerves during stimulation of the mesothoracic nerve 2. In all examples, responses decreased either in intensity and duration, or disappeared completely after SEG removal.

## 3 Results

### 3.1 Various proprioceptive inputs are processed by the SEG to elicit motor responses

Our recently reported results (Knebel et al. 2017b) suggest a role of the SEG in modulating the coupling between segmental leg CPGs in the absence of leg sensory feedback. Here we sought to uncover the SEG modulation of sensory-motor responses.

Motor responses of the middle leg of an intact locust were monitored following mechanical movement of the ipsilateral hind leg, and electrical stimulations of its main sensory nerve, Nb5. The latter includes axons of hair sensilla, campaniform cells, and chordotonal organs from the leg. The responses included both multi-joint middle-leg movements and fully coordinated steps. The probability of response was compared among intact, brainless and headless (brain and SEG removal) animals, showing similar response profiles for the mechanical and the electrical stimulations (N=19; Fig 1A). Response probabilities remained similar after decerebration, but were significantly reduced after full decapitation (chi test with Bonferroni correction for three comparisons, p<0.001 between intact and decapitated animals and p<0.01 between decerebrated and decapitated animals). These results indicate that the SEG plays a role in the induction of sensory-motor responses among the legs.

We continued testing the influence of the SEG on leg sensory-motor responses in reduced preparations of the isolated SEG and the thoracic ganglia chain. The responses of different leg motor circuits to different sensory-like stimulations were monitored. We found various stimulation protocols that elicited inter- and intrasegmental response in leg motor nerves, which were affected by SEG inputs. Fig. 1A & B present two examples of stimulation of N5b in both the meso- and metathoracic ganglion. As can be seen, induced motor responses underwent some change following SEG removal, in accord with the *in-vivo* experiments (Fig. 1A). Similarly, stimulation of the mesothoracic N2, which among others innervates thoracic hair sensilla and chordotonal organs (Bräunig et al. 1981), resulted in an intersegmental coordinated response, which was again dependent on SEG inputs (example in Figure 1C). In the different utilized stimulation protocols, some trials were observed to result in bursting activity that lasted for a few seconds following the stimulation. This activity alternated between adjacent legs: a coordinated pattern that can be described as fictive stepping (Figure 1A & C). We focused the rest of the investigation on N5b stimulation protocol *in-vitro*.

### 3.2 The SEG induces antiphase intersegmental sensory-motor responses

To further characterize the response to the metathoracic N5b stimulation, we monitored the activity at the ipsilateral meta- and mesothoracic depressors in 19 preparations. Each experiment comprised three stimulations with intact SEG, with an interval of 30 sec between them, and similarly three stimulations following removal of the SEG (pictogram in Fig. 2A). A consistent reduction in average spike frequency in the metathoracic motor output was almost always observed following stimulation, regardless of the presence of the SEG (red lines, Figure 2A; Wilcoxon signed-rank test with Bonferroni correction for six comparisons, p<0.001). In contrast, a significant increase in spike frequency was seen for the ipsilateral mesothoracic depressor nerve when the SEG was intact (blue lines in Figure 2B, left trace; Wilcoxon signed-rank test with Bonferroni correction for six comparisons, p<0.01). Interestingly, this intersegmental effect was abolished following removal of the SEG, after which no consistent response to the stimulation was observed (a mixture of blue and red lines, Fig. 2B right trace).

**Fig. 2.**
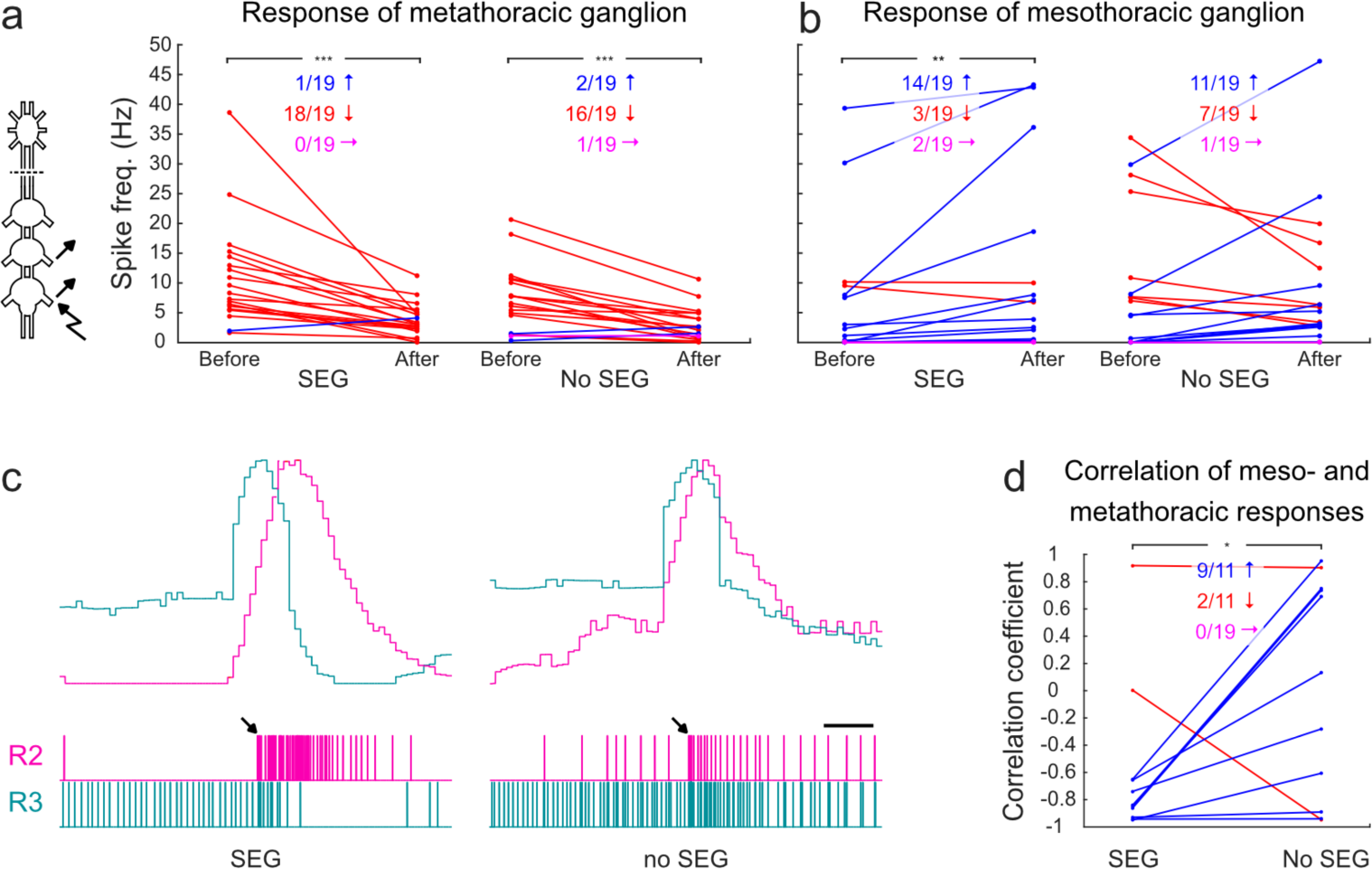
Analysis of inter- and intra-segmental responses to metathoracic N5b stimulations, with and without the SEG. (A) Average spike frequencies of the metathoracic depressor nerve before and after the stimulation, with and without the SEG. Blue, red, and magenta lines represent elicited, or decreased activity, or no change following a stimulation, respectively. The matching color numbers indicate the proportions of the experiments in each of these response categories. Significant changes in motor activity at the same segment were found both with and without the SEG. In all panels, analysis was made for a time window of 4 sec before and 4 sec after stimulation. (B) As in (A), but for the mesothioacic depressor ipsilateral to the stimulation. Here significant responses were found only when the SEG was intact. (C) Activity profiles were calculated to determine cross-correlation between responses at the ipsilateral segments. For the analysis, we used the vectors of activity densities of the meso- and metathoracic depressors (R2 and R3 and the corresponding red and blue curves), calculated in 100 msec time bins. Arrows indicate stimulation onsets. Scale bar is 1 sec. (D) Correlation coefficient values, calculated from the density vector of 4 sec following the stimulation onset, reveal an anti-phase correlation for most of the intact SEG preparations vs. highly variable responses for the non-SEG ones.

To examine the influence of the SEG on the overall intersegmental activity we calculated the correlation between the meso- and metathoracic activity profiles following the stimulation. We characterized the temporal properties of the activity by calculating correlation coefficients between vectors of activity densities (curves in Figure 2C). In accordance with the results of averaged activity changes (Figure 2A & B), and taking into account the temporal properties of the ipsilateral segment responses, an anti-phase correlation was observed for most preparations with an intact SEG (Figure 2D). This sensory-induced alternating activity resembled the coordination pattern seen in normal walking gaits. Following removal of the SEG, the correlation markedly changed: the anti-phase activity disappeared, and the overall post-stimulation activity became quite synchronized (Fig. 2D; Wilcoxon signed-rank test, p<0.05).

### 3.3 SEG sensory-related inputs descend ipsilaterally

Our findings suggested a direct involvement of the SEG in sensory-motor processing of intersegmental information transfer within the thoracic ganglia. To track the SEG descending activity in response to our sensory-like stimulations, we monitored the inter-ganglia connectives ipsilateral to the stimulation site. SEG descending interneurons (DINs) could be identified by overlaying traces recorded simultaneously from the SEG-prothoracic, and pro-mesothoracic connectives (example in Figure 3A and analysis in 3B and 3C). An example of the activity of a SEG DIN following metathoracic N5b stimulation is shown in Figure 3, displaying both activity density and raster plots of the individual stimulations and their average. A strong activation of DINs can be seen following the sensory stimulations, indicating an involvement of the SEG descending inputs in the sensory-induced motor responses, conveyed via ipsilateral tracts to the stimulated side. In accordance, the rhythmic footprint of the DIN activity (Fig. 3C bottom trace) could correspond with rhythmic motor output to the legs.

**Fig. 3.**
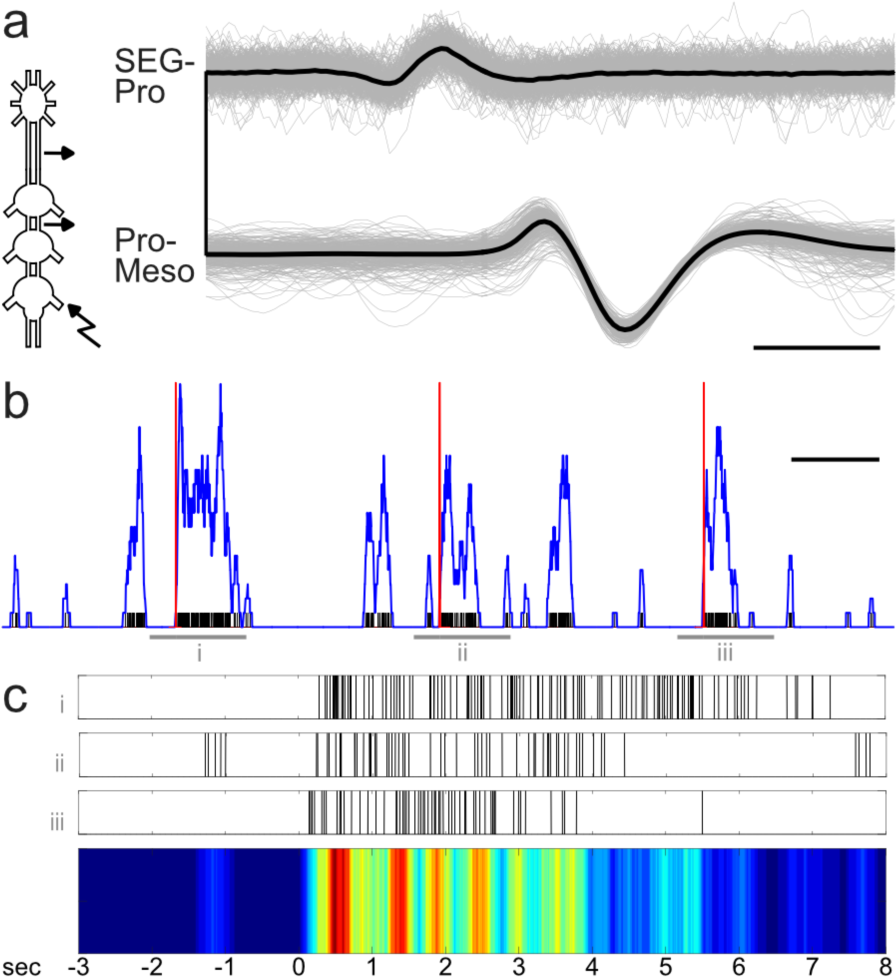
An example of a SEG DIN becoming active following the metathoracic 5b stimulations. (A) An overlay of the SEG-prothoracic and pro-mesothoracic connective recordings, fixed to the firing of one neuron. The consistent delay in recorded spike between the electrodes (~2 ms) fits axonal conduction speed for the distance between the electrodes, and indicates that it is a SEG DIN activity. Scale bar is 2 ms (B) The on times of the spikes (black) are presented on top of the on times of the stimulations (red). The blue line represents a smoothing of the on times vector of the spikes. Scale bar is 10 sec. (C) The on times of the DIN firing before and after the three stimulations (three upper traces, numbered by their corresponding time windows shown in (B)), and their color-coded summation (bottom trace). Zero time indicates stimulation onset. Color index indicates the level of activity (normalized units).

### 3.4 Sensory information ascends contralaterally

About 70% of the SEG DINs project contralaterally to the soma (Kien et al. 1990; Gal and Libersat 2006). Therefore, if the descending SEG commands following sensory-like stimulation are transferred ipsilaterally to the leg motor centers, as our results suggest, there is a high probability that the ascending sensory information is received on the SEG side contralateral to the stimulation. To test this hypothesis, we repeated the metathoracic N5b stimulation protocol in preparations in which the major connectives were sectioned at different locations, while recording the mesothoracic depressor ipsilateral to the stimulation side. First, the meso-metathoracic connective ipsilateral to the stimulation side was cut (pictogram in Figure 4A). Both before the cut and following it, the mesothoracic depressor responded in excitation to the stimulation (Figure 4A-B; N=12, Wilcoxon signed-rank test with Bonferroni correction for nine comparisons, p<0.001 and p<0.005, respectively). As before, following the SEG removal the response disappeared (Wilcoxon signed-rank test with Bonferroni correction for nine comparisons, p<0.001). Next, we cut the meso-metathoracic connective contralateral to the stimulation site (pictogram in Figure 4C) and observed that, as a result, the response disappeared (Figure 4C-D; N=12, Wilcoxon signed-rank test with Bonferroni correction for nine comparisons, p<0.001 and no significance, respectively). As expected, following the SEG removal, there was still no response.

**Fig. 4.**
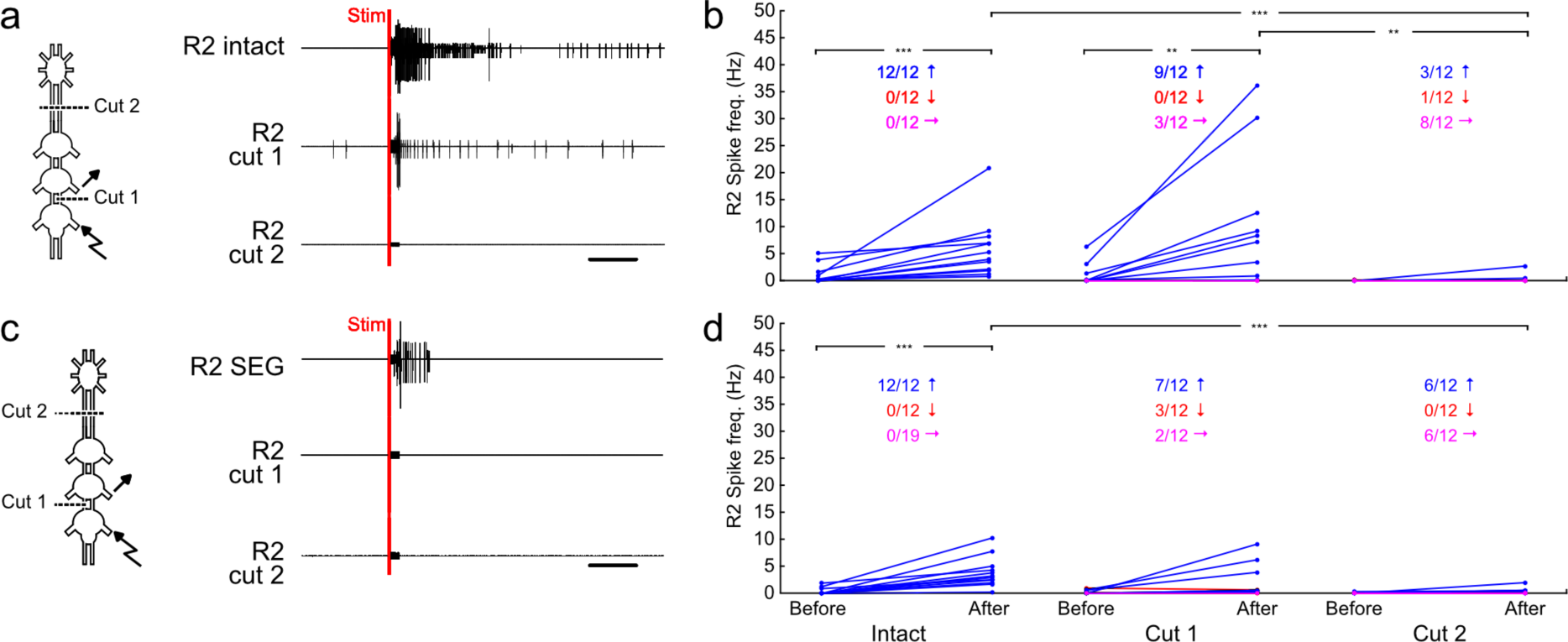
Specific lesioning experiments reveal the pathway of transferring the sensory signals. (A) In the first set of experiments, cuts were made at the ipsilateral connective between the stimulated meta- and recorded mesothoarcaic segments (cut 1) and then at the two connectives between the SEG and the prothoracic ganglion (cut 2). An example of the corresponding response to stimulations, before cutting (top trace), after cut 1 (middle trace), and after cut 2 (bottom trace) is plotted. Scale bar is 1 sec (B) Spike frequencies of the ipsilateral mesothoracic depressor before and after stimulation. Blue, red, and magenta lines represent elicited, or decreased activity, or no change following a stimulation, respectively. The matching color numbers indicate the proportions of the experiments in each of these response categories. Overall, the data indicate that responses remained after cutting the ipsilateral connective, but were abolished following SEG-removal. Statistical analysis of the activity after stimulation reveals a difference between SEG and No SEG group but not between the SEG and the ipso-cut ones. (C) and (D) as in (A) and (B) but for the second experimental set, with cuts of the contralateral connective between the meso- and the meta-thoracic ganglia (cut 1) and between the SEG and the prothoracic (cut 2). Here, response to stimulations were abolished already after cut 1, suggesting that the response is mediated by ascending pathways through the contralateral connective.

Double staining, in two preparations, of the metathoracic N5b and the contralateral connective (Pictogram in Figure 5a), has revealed several ascending neurons whose neurites project contralaterally to their soma and overlap with dendritic fibers of the local N5b sensory projections (example in Figure 5B-D). Although these cannot indicate that synaptic connections between the two exist, they propose a possible route for transferring the proprioceptive information to the SEG via ascending neurons that project contralaterally to their soma. First, the N5b sensory projections were found to be present only locally within the metathoracic hemiganglion, neither projecting to the contralateral side nor to more rostral segments (N=2). Therefore, the information transfer to the SEG, and to other hemiganglia must be conveyed by interneurons. Second, the overlap between the sensory projections and ascending interneurons opens the possibility that the sensory information ascends through monosynaptic connections.

**Fig. 5:**
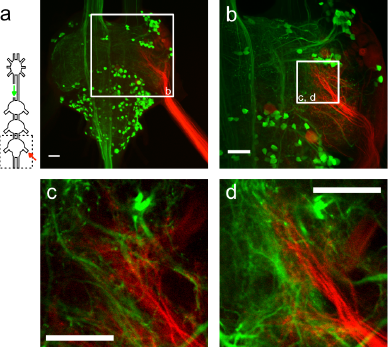
Double backfill staining of the left connective, between the SEG and the prothoracic ganglion and the metathoracic right N5b. (A) Maximum intensity Z-projection of the entire ganglion. (B) same as (A) but for the region marked by a white square in (A). (C) and (D) single slices from the confocal scan of the area marked in (B) at different depths of the ganglion show proximate arborization of sensory projections and ascending neurites. Scale bars are 100 µm.

## 4 Discussion and Conclusions

While various studies have demonstrated that insect locomotion is dependent on inputs from the intact SEG (e.g. Kien 1983; Gal and Libersat 2008), little is known about the interactions of the SEG with the different leg motor centers. Here we have shown that inputs from the locust SEG modulate interactions between sensory inputs and motor output. Consistent with our previously suggested roles of the insect SEG in leg control (David et al. 2016; Knebel et al. 2017b), we have shown here that SEG descending signals facilitate intersegmental transfer of sensory information for a functional coordination.

### 4.1 Interactions of sensory inputs and higher motor centers in walking behavior

Sensory inputs are instrumental for generating functional gaits by insect pattern- generating networks (stick insect: Borgmann et al. 2007, 2009; Daun-Gruhn 2011; locust: Runion and Usherwood 1968; Newland and Emptage 1996). Based on our knowledge of sensory modulation of other rhythmic behaviors (e.g. lobster STG; for review: Marder and Bucher 2007), this effect is probably manifested by way of multi-targeted modulation, including that of the pattern-generating networks themselves, as well as their interconnections. Sensory signals should have a behavioral-context-dependent effect: i.e., in different contexts the same inputs may be required to generate different effects. This can be achieved via interactions with descending inputs from higher motor centers (for a review on vertebrates: Hultborn 2001). For example, Mu and Ritzmann (2008) demonstrated an ability of (unspecified) descending inputs from the cockroach head ganglia to modulate leg reflexes, and even to reverse their motor output.

It should be noted that insect leg sensory nerves deliver a rich and complex sensory signal, originating in different sensory receptors (Bräunig et al. 1981; Ayali et al. 2015b). It was out of the scope of the current study to characterize various sensory signals. Rather we have used mechanical leg stimulation or electrical nerve stimulation for the purpose of testing for a role of the SEG in the control of intra- and intersegmental motor responses to sensory perturbation. The similar results obtained in the different stimulation protocols utilized allowed us to further examine the interactions between the SEG and the sensory-motor pathways in an *in vitro* preparation.

Our results indicate that the locust SEG plays a significant role in intersegmental, but not intra-segmental, transfer of leg sensory-related information. This differential activity, connecting different local leg motor centers, suggests that the SEG coordinates the overall output to the legs in response to local sensory inputs. The long time scale and high intensity of the motor responses to the stimulations possibly indicate the participation of neuromodulatory substances that could be secreted by SEG cells. It has been previously shown that the activation of leg-related octopaminergic dorsal unpaired median neurons (DUM3,4,5) in the metathoracic ganglion resulting from sensory stimulation (of tactile hairs and campaniform sensilla) was abolished following disconnection of the head ganglia (Field et al. 2008), and specifically of the SEG (see also in Manduca: Pflüger et al. 1993; Johnston et al. 1999). We have also recently reported that SEG DUM neurons receive an efference copy of the legs’ motor output and deliver descending commands (Knebel et al. 2017b). The long and highly variable response latencies to the different stimuli observed in the current study, with median delays of up to 480 ms, support the involvement of multi-synaptic ‘sensory to neuromodulatory’ pathways. These time scales are in accord with the observed prolonged responses in the presence of SEG descending inputs. It is likely, however, that these pathways operate in parallel in directing afferent signals to motor connections.

In both of our previous recent reports (Knebel et al. 2017a,b) we were unable to produce walking-like rhythms *in vitro* (Figure 6A). Moreover, we reported that the ipsilateral leg CPGs are probably wired to oscillate in phase (Knebel et al. 2017a). We therefore assumed that in order to coordinate a functional gait, sensory information is required. The results presented here indicate that sensory information, together with descending commands, can break the observed ipsilateral CPG in-phase coupling, and induce walking-like alternations among adjacent legs even *in vitro* (Fig. 2D). This further demonstrates the great importance of sensory information for adequate walking behavior. This may be in contrast to other locomotory behaviors, such as swimming or crawling, that take place in more homogenous environments (for a review: Büschges et al. 2011).

**Fig. 6:**
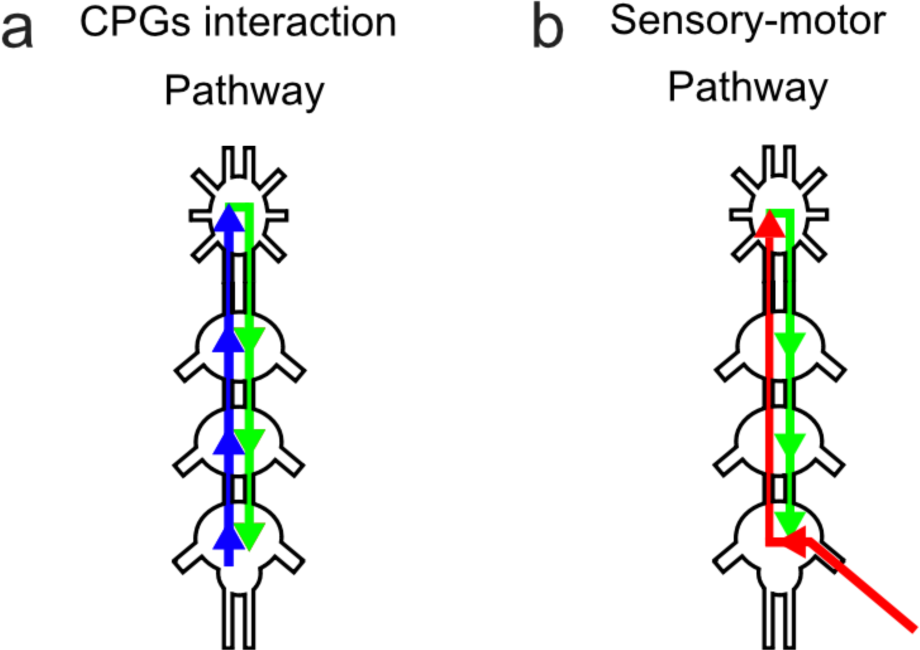
Schematic maps of the motor and sensory-motor pathways. (A) Schematic map of the suggested predominant connections among the leg-CPGs, based on Knebel et al., (2017a,b). The blue line represents the information transfer among the CPGs and to the SEG, while the green line depicts the descending information from the SEG to the leg CPGs. (B) Schematic map of the suggested pathways of sensory-motor information transfer. The red line represents the sensory nerve stimulation and its path up to the SEG, while the green line depicts the descending commands from the SEG to the leg motor centers. Note that the predominant connections among the legs CPGs are ipsilateral with a bilateral SEG connection, and that sensory information ascends contralaterally to the stimulated nerve, and crosses the midline back in the SEG to affect the ipsilateral legs.

As described in the methods, it should be noted that the stimulation was not attempting to activate a specific proprioceptive organ, but to generate a general sensory perturbation. In addition, we note that findings obtained *in vitro* are not conclusively applicable to the complete behavior of the intact animal. However, as has often been discussed (e.g. Fuchs et al. 2012; David et al. 2016), they do encompass valuable information on how behavior is generated from (or despite) the hardwired circuitry, which is still intact in the isolated preparation.

### 4.2 Mapping of the information transfer pathways

Previous reports have shown that while brain-less insects walk normally, even more readily than intact animals (e.g. Kien 1983; Gal and Libersat 2006), SEG-less animals do not engage in spontaneous walking. It is therefore possible that SEG descending inputs provide some general excitation to local walking centers and, consequently, that their activation, either by sensory feedback or descending brain commands, would require less synaptic drive. The SEG-dependent intersegmental responses to sensory-like stimulation reported here might be the result of such a general excitation. However, other evidence suggests differently. Knebel et al. (2017b), observed no change in the excitability of the leg CPGs with and without inputs from the SEG. Similarly, in the current study no significant changes in activity were found between the experimental conditions before stimulation (with and without the SEG; Figure 2 & 4). Furthermore, we have demonstrated here that sensory-stimulation activated SEG DINs irrespective of and beyond any basal excitation of the leg motor centers by the SEG. Finally, if the observed intersegmental response to stimulation was mediated merely by general excitation from the SEG, the responses should have been insensitive to our differential connective cuts (Figure 4). Therefore, we suggest that the SEG receives specific sensory information, processes it, and delivers sensory-induced commands to the legs.

Further work is required to completely map the anatomical and physiological interconnections between leg sensory structures, the thoracic CPGs, and the SEG (following Bräuning et al. 1981). However, based on our neuroanatomical staining, and unlike other sensory projections (e.g. mesothoracic N2; Bräunig et al. 1981), the metathoracic N5b does not appear to project to the other thoracic or head ganglia (Figure 5A). Namely: our staining results do not support direct connections between the main leg sensory organs and the SEG. The intersegmental sensory motor response presented here is, therefore, conveyed through ascending and descending interneurons.

Based on the metathoracic N5b backfill staining, we conclude that sensory information from the leg is received in the proximate hemiganglia (Figure 5A). As our physiological findings from lesioning the different pathways suggest, the information then passes to the contralateral side within the ganglion and ascends to the SEG (Figure 4). It is possible that connections between the sensory organs and ascending interneurons are monosynaptic, and as the double backfill staining indicates, a few ascending interneurons indeed send their neurites contralaterally to their soma and share similar arborization sites with N5b sensory projections (Figure 5C-D). The ascending information is received in the SEG on the contralateral side to the leg, and as most SEG DINs project contralaterally to their soma, their descending signals will be delivered to the thoracic segments through the connective ipsilateral to the stimulated leg (Figure 3). This suggested map is schemed in Figure 6B.

### 4.3 Comparison to vertebrates

Our knowledge of the higher motor centers in vertebrates (mostly mammals) suggests a clear functional divergence between the motor cortex and basal ganglia, both of which are involved in the planning, control, and execution of voluntary movements, and the brainstem, which is more closely associated with spinal motor circuits. In insects too, two neuronal centers within the head ganglia have been attributed a role in motor control: the brain central complex (e.g. Bender et al. 2010; Martin et al. 2015) and the SEG. While the former was suggested to modify the overall walking behavior in regard to environmental requirements, the latter is embryonically related to the thoracic ganglia and functionally associated with their CPGs. Of special interest is the possible homology (or perhaps even analogy) between the higher motor centers in vertebrates and those of insects. Recent studies have compared the development and function of the insect central complex and the vertebrate basal ganglia (Wessnitzer and Webb 2006; Strausfeld and Hirth 2013). Our understanding of the role of the SEG in insect locomotion, not to mention its evolutionary similarities with the vertebrate brainstem, is much more limited. However, the current findings support functional and anatomical parallels between the SEG and the brainstem, as previously suggested by Schoofs et al. (2014). The suggested intersegmental contralateral transfer of sensory signals via the SEG, as elaborated above and in Figure 6B, is in accord with that in the sensory-ascending pathways in vertebrates. These decussate to the contralateral side, either close to the entrance of the corresponding primary afferents, as is the case for pain or temperature pathways, or at the brainstem, as for tactile and proprioception signals (for a review: Guertin 2013). In both pathways, the sensory neuron forms an ipsilateral synapse on an ascending interneuron, and the latter crosses the midline of the system to supply the contralateral hemisphere. As in the vertebrate brainstem, anatomical observations suggest that the majority of SEG-DINs possess contralateral axons (Kien et al. 1990; Gal and Libersat 2006).

### 4.4 Conclusions

By recording the motor responses to leg sensory stimulations we show that inputs from the locust SEG modulate interactions between legs’ sensory inputs and motor output. The SEG is essential for the intersegmental, but not for the intrasegmental transfer of sensory signals. Specific sections along the intersegmental connectives indicate that the sensory inputs are mainly transferred to the SEG via the contralateral tracks, similar to ascending pathways in vertebrates.

## Acknowledgments

This work was supported by the Young Scholar Fund, University of Konstanz, the Lion foundation for a travel scholarship to JW and also in its final stages by the German Research Council (DFG; Grant RI 2728/2-1). We thank Prof. Hans-Joachim Pflüger and Prof. Peter Bräunig for productive discussion and support.

